# The connecting cilium inner scaffold provides a structural foundation to maintain photoreceptor integrity

**DOI:** 10.1101/2021.08.27.457921

**Authors:** Olivier Mercey, Corinne Kostic, Eloïse Bertiaux, Alexia Giroud, Yashar Sadian, Ning Chang, Yvan Arsenijevic, Paul Guichard, Virginie Hamel

## Abstract

Retinal degeneration is a leading cause of human blindness due to progressive loss of ciliated photoreceptors cells. While this degradation can be associated with cohesion defects of the microtubule-based connecting cilium (CC) structure, the underlying mechanism is not understood. Here, using expansion microscopy and electron microscopy, we reveal the molecular architecture of the CC and demonstrate that microtubules are linked together by a CC-inner scaffold (CC-IS) containing POC5, CENTRIN and FAM161A. Monitoring CC-IS assembly during photoreceptor development in mouse reveals that it acts as a structural zipper, progressively bridging microtubule doublets and straightening the CC. Consistently, *Fam161a* mutations lead to a specific CC-IS loss and trigger microtubule doublets spreading, prior to outer segment collapse and photoreceptor degeneration, providing a molecular mechanism for *retinitis pigmentosa* disease.

**One Sentence Summary:** The connecting cilium inner scaffold acts as a structural zipper granting photoreceptor integrity.

## Main Text

The retina is a thin tissue lining the back of the eyeball containing photoreceptor cells, which convert light inputs into electrical signals, a process crucial for the detection of visual stimuli. These highly specialized ciliated cells are partitioned into two main regions, a photosensitive modified primary cilium called outer segment (OS) and a cell body named inner segment, both connected via a thin bridging structure known as connecting cilium (CC). This structural linker, emanating from a mature centriole, is made of nine microtubule doublets (MTDs) that extend distally to form the OS onto which hundreds of stacked membrane discs containing phototransduction proteins are positioned (*1*–*3*). Mutations in the microtubule binding protein FAM161A, found to localize at the CC, have been associated with the human pathology *retinitis pigmentosa 28* (RP28), a subtype of the most prevalent human inherited retinal disease (*4*–*11*) with an incidence of 1/4000 worldwide. Mouse models of FAM161A-associated RP28 have revealed structural defects in the CC displaying spread MTDs and disturbed membrane discs organization that underlined photoreceptor loss (*9, 12*). Similarly, mutations in the CC-localized proteins POC5, CENTRIN and POC1B have all been associated to retinal pathologies displaying photoreceptor degeneration (*13*–*15*).

Besides being present at the CC, these four proteins are located at the level of centrioles, and compose the recently identified inner scaffold (IS) structure connecting neighboring microtubules blades (*16*). Depletion of IS components leads to centriole architectural defects, hinting for a role of this structure in the structural cohesion to the entire organelle (*16, 17*). These four IS proteins being also present at the CC, we hypothesized that a similar IS structure might exist at the level of the CC to provide the structural cohesion of the MTDs, thus granting OS integrity.

To test this hypothesis, we set out to reveal the molecular architecture of the CC by using and optimizing Ultrastructure Expansion Microscopy (U-ExM) (*18*) on adult mouse retinal tissue (**Fig. S1A**). To first validate it, we analyzed at low magnification the overall organization of the tissue after expansion and demonstrated the preservation of the different cell layers of the retina, notably the photoreceptors cells (**Fig. 1A, B**). Moreover, staining photoreceptor cells for *α*/β-tubulin revealed microtubule structures at higher magnification, thus allowing the visualization of both mother and daughter centrioles, the CC and the extending OS axoneme, which forms an enlarged region immediately after the CC, that we named the bulge (**Fig. 1B, S1B**). Additionally, we examined the localization of the centriolar IS component POC5 and consistently found it in the central core region of centrioles and also clearly decorating the entire CC region as previously reported (*15, 16*) (**Fig. 1B)**. From this result, we next determined the expansion factor resulting from this optimized protocol using photoreceptor centriole diameter as a ruler and warrant of structural integrity. By comparing it to measurements of U2OS human centrioles, we found an expansion factor of 4.2 on average (**Fig S1C-F**), matching the previously published values, thus validating our approach (*16, 19*).

**Fig. 1.**
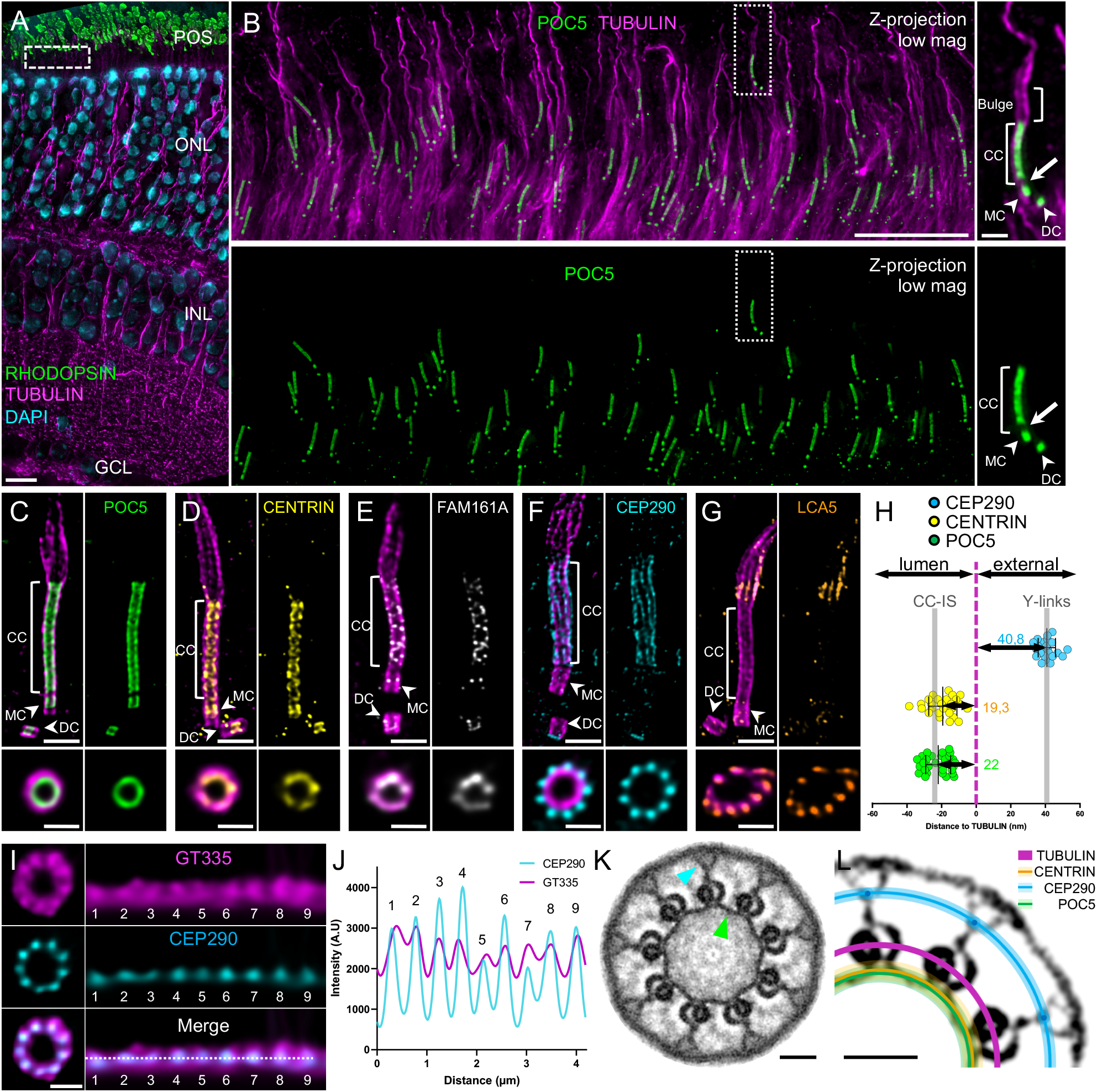
Molecular mapping of the mammalian photoreceptor connecting cilium. (**A**) Low magnification U-ExM image of a P14 mouse retina highlighting the preservation of the different retina layers. Note that the RPE layer is removed with the dissection. Scale bar: 10 µm. (**B**) Expanded P14 photoreceptor layer (equivalent region to the white dashed line depicted in (**A**)). Inset shows details of one photoreceptor cell. Arrowheads indicate centrioles. Arrow depicts the gap between centriole and CC POC5 signals. Scale bar: 5 µm (inset: 500 nm). (**C-G**) Confocal U-ExM images of photoreceptors stained for tubulin (magenta) and POC5 (**C**: Green) or CENTRIN (**D**: yellow), FAM161A (**E**: gray), CEP290 (**F**: cyan) and LCA5 (**G**: orange). Lower panels show transversal views of the CC for each staining. Arrowheads indicate centrioles. Scale bars: side view= 500 nm; transversal view= 200 nm. (**H**) Distances between the maximum intensity of POC5 (green), CENTRIN (dark yellow) and CEP290 (cyan) compared to tubulin (magenta) calculated from transversal view images. Grey bars indicate the values obtained from the simulation in Fig. S3. n≥ 2 animals per staining. See Table S1. **(I**) Transversal view (left) and polar transform (right) of GT335 (magenta) and CEP290 (cyan) signals revealing overlapping 9-fold symmetry. Scale bar: 200 nm. (**J**) Plot profiles of CEP290 (cyan) and GT335 (magenta) polar transform of (I). (**K**) Symmetrized EM image of a P14 CC transversal section revealing an inner ring decorating MTDs (green arrowhead) and Y-links bridging MTDs to the membrane (blue arrowhead). Scale bar: 50 nm. (**L**) Model representing relative positions calculated in (H) of POC5 (green line), CENTRIN (dark yellow line), CEP290 (cyan dot and line) to tubulin (magenta) on a contrasted symmetrized EM picture of a CC. Light color lines represent the SD for each protein. Scale bar: 50 nm. POS= Photoreceptor Outer Segments; ONL= Outer Nuclear Layer; IPL= Inner Nuclear Layer; GCL= Ganglion Cells Layer; MC= mother centriole; DC= daughter centriole; CC= connecting cilium.

We next investigated the precise localization of the IS components POC5, CENTRIN and FAM161A, with the exception of POC1B owing to the lack of appropriate antibodies. We also inspected the distribution of the protein CEP290, known to localize along the CC in super-resolution microscopy (*20*), and Lebercilin (LCA5), a proposed FAM161A interactor (*10*), whose mutations have been linked to Leber congenital amaurosis, a retinopathy causing severe visual deficiency from the first year after birth (*21, 22*). Consistently with our previous work, all three IS proteins were found in the central core region of centrioles, where the IS structure lies (*16*) (**Fig. 1C-E**). In addition to confirming POC5, CENTRIN and FAM161A localization along the CC, we could observe that these proteins are lining the inner part of the microtubule wall, although FAM161A antibody displayed a weaker and dotty signal probably due to partial epitope conservation from mouse to human (**Fig. 1C-E**). We also noted a gap between protein signals in the centriole and the CC, suggesting that these two regions are independent (**Fig. 1B**, white arrow). In contrast, we found that CEP290, capping the distal end of daughter centrioles, decorates the external part of the microtubules along the CC with a 9-fold symmetry revealed from transversal sections (**Fig. 1F**), and that most of LCA5 lined MTDs in the bulge region, occasionally being weakly present at centrioles and along the CC. While LCA5 has been mostly described along CC, this localization corroborates previous immunogold results (*22*) (**Fig. 1G**).

Next, to precisely map these different proteins relative to the MTDs within the ultrastructure of the CC, we measured the distance between peak intensities of POC5, CENTRIN and CEP290 compared to tubulin from transversal section images, omitting FAM161A as the quality of the signal did not allow precise measurements. We found that POC5 and CENTRIN are internal to the microtubule wall by **∼**20 nm similarly to its position on the IS at centrioles (*16*), while CEP290 is **∼**41 nm external to the MTDs (**Fig. 1H**). By co-staining CEP290 with the external tubulin glutamylation marker GT335, we also unveiled that CEP290 is contiguous to the MTDs (**Fig. 1I, J**). To correlate these measurements with the CC ultrastructure, we then performed transversal EM sections of wild-type P14 mouse photoreceptors (**Fig. 1K** and **Fig. S2**). By applying a nine-fold symmetrization to increase the contrast, we revealed the presence of the Y-links structures connecting the MTDs to the ciliary membrane as well as an inner ring connecting neighboring MTDs, previously described as a transition zone ring (*23, 24*) and reminiscent of the centriolar IS, which we therefore dubbed CC-inner scaffold (CC-IS) (**Fig. 1K, Fig. S2**). By superimposing the measured distances of POC5, CENTRIN and CEP290 onto a symmetrized EM image, we demonstrated that POC5 and CENTRIN perfectly co-localize with the CC-IS and that CEP290 localizes at the level of the Y-links region (**Fig. 1L**). To ascertain these results, we performed an U-ExM simulation from EM symmetrized images (**Fig. S3**) and confirmed that these structural elements depict similar localization signals as the mapped proteins. Taken together, these molecular and ultrastructural analyses confirmed the existence of an inner scaffold at the CC, linking MTDs together.

We then investigated how and when the CC, and notably the CC-IS, assembles during early postnatal development of photoreceptors in mouse, between days 4 and 60 (P4 and P60). We first monitored rod OS maturation at low magnification using tubulin and RHODOPSIN, the sensory protein that captures light and promotes conversion into electric signal (*25*). We found that U-ExM recapitulated OS formation with the appearance of discrete RHODOPSIN signals between P4 and P7 that gradually extend to form mature OS at P30-P60 (*3*) (**Fig. 2A**). Then, we analyzed at higher magnification the CC-IS formation using POC5 signal as a proxy (**Fig. 2B**). We found that at P4, while all centrioles are POC5-positive and CC microtubules are already elongating, only 42% of photoreceptors displayed a POC5 staining at the CC (**Fig. 2B, K**). Early after, at P7, POC5 is detected in 95% of the growing CC, spanning on average 415 nm (**Fig. 2B, E and K**). POC5 signal then elongates in parallel to OS maturation until it reaches its final length of around 1500 nm at P60, similar to the described length of a mature CC (*26*) (**Fig. 2B, E**). By measuring the relative position of the proximal and distal POC5 signal in the CC compared to the centriolar distal end, we also noticed that POC5 initially appears randomly along the CC, away from the mother centriole, and that its coverage elongates towards both the mature centriole and the distal end of the CC, suggesting that the CC-IS structure growth is bidirectional (**Fig. 2B, H, J, L, Fig. S4A**).

**Fig. 2.**
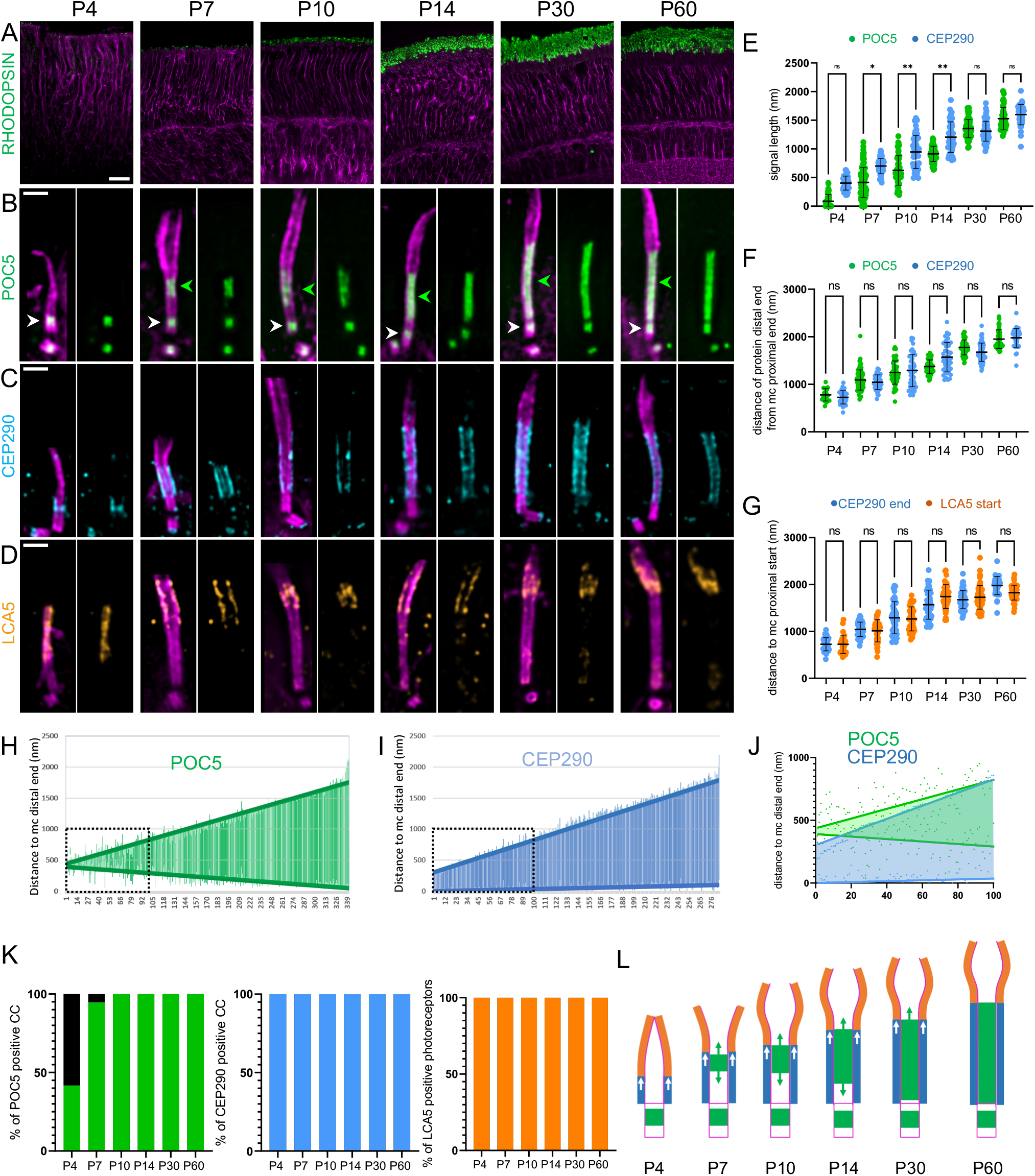
Photoreceptor CC-inner scaffold assembly. (**A**) Low magnification of expanded retinas showing OS formation from P4 to P60 stained with RHODOPSIN (green) and tubulin (magenta). Scale bar: 20 µm. (**B-D)** Expanded photoreceptors stained for tubulin (magenta) and POC5 (green, B), CEP290 (cyan, C) or LCA5 (orange, D) from P4 to P60. Note that CEP290 is also capping the daughter centriole. Green arrowheads= CC-IS; white arrowheads= centriole IS. Scale bar: 500 nm. (**E**) Quantification of POC5 (green) and CEP290 (cyan) signal length over time. ≥ 3 animals per time point. (**F**) Quantification of the distance of POC5 (green) and CEP290 (cyan) signal distal ends in the CC to the mother centriole proximal end from P4 to P60. ≥ 3 animals per time point. (**G**) Distance of CEP290 signal distal end (cyan) and LCA5 signal start (orange) to the mother centriole proximal end. ≥3 animals per time point. Note that CEP290 measurements are the same as in (F). (**H, I**) Length and position of POC5 (H: green) and CEP290 (I: cyan) signal within the CC. Note that signals are sorted by length and distances and compared to MC distal end. Thick green (POC5) and cyan (CEP290) lines represent linear regression curves. Black dashed square represents the inset highlighted in (J). ≥3 animals per time point. (**J**) Comparison of the 100 shortest CC reveals POC5 (green) bidirectional and CEP290 (cyan) unidirectional growth, respectively. (**K**) Percentage of CC positive for POC5 (green), CEP290 (cyan) and LCA5 (dark yellow) from P4 to P60. ≥3 animals per time point. (**L**) Model showing inner scaffold (green) and CEP290 (cyan) growth over time, dictating the final length of the CC and the starting point of the bulge region (orange). Means, percentage and standard deviations are listed in Table S1.

We further investigated whether the CC-IS assembly coincides with that of the Y-links and the bulge region defined by CEP290 and LCA5, respectively (**Fig. 2C-G**). We found that both CEP290 and LCA5 appear earlier than POC5 from P4 onwards as 100% of photoreceptors displayed a signal for these two proteins (**Fig. 2K**). In contrast to the bidirectional growth of POC5, CEP290 extends directly from the end of the underlying mother centriole and exhibits a unidirectional growth, indicating that the assembly of these two structural modules is probably independent (**Fig. 2C, E, I, J, L and Fig. S4B**). However, we noticed that POC5 and CEP290 signal distal ends grow concomitantly and correspond to the starting point of LCA5 signal (**Fig. 2F, G**). This observation suggests that the growth of the CC region, encompassing the bidirectional elongation of the CC-IS and unidirectional Y-links growth, displace distally the beginning of the bulge where membrane discs are formed (**Fig 2C-G, L**).

A key prediction of the CC-IS function is that it bridges and maintains neighboring MTDs cohesion during its elongation. To test this hypothesis, we next explored the MTDs diameter during CC formation. Indeed, while monitoring the growth of the CC-IS, we observed that the axoneme diameter seems wider in the absence of POC5, compatible with the fact that the absence of the CC-IS coincides with spread MTDs **(Fig. 3A)**. To confirm this hypothesis, we measured between P4 and P60 the tubulin diameter at different positions along the CC photoreceptors: at the end of POC5 signal (0), but also 150 nm below (−150) and above (+150) (**Fig. 3B**). At every time point, we found a significant tubulin diameter enlargement distally to the end of the POC5 signal, suggesting that CC-IS appearance gathers MTDs together early and throughout OS development (**Fig. 3B**). Consistently, we found that the position of the microtubule enlargement corresponds to the distal end position of the CC-IS marked by POC5 (**Fig. 3C, D**). To test whether this microtubule spread is correlated with the absence of the CC-IS at the ultrastructural level, we next analyzed EM transversal sections of P14 photoreceptors (**Fig. 3E**). As expected, we found that the bulge region, identifiable by the absence of Y-links and displaying randomly shaped associated membranes, is lacking the CC-IS (**Fig. 3E**). Moreover, this region has a larger perimeter and a less circular shape, correlating the lack of cohesion between MTDs (**Fig. 3F, G**). The absence of microtubule cohesion was even more pronounced in the distal cilium region where singlet microtubules can be observed (**Fig. S1B**). These results confirmed that the growing CC-IS gathers neighboring MTDs during elongation, acting as a structural zipper, during CC assembly.

**Fig. 3.**
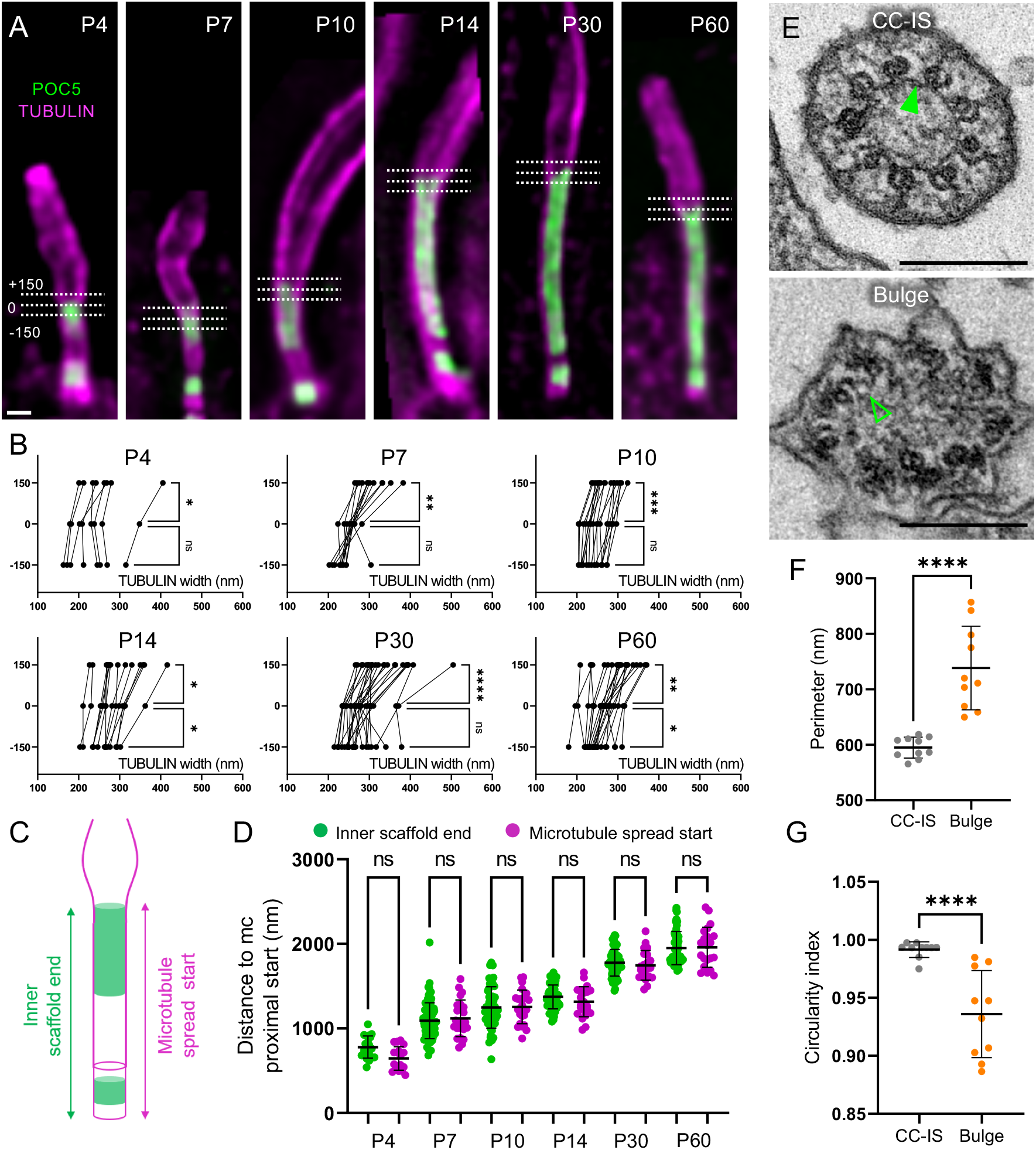
The CC-inner scaffold acts as a structural zipper maintaining MTDs cohesion. (**A**) Expanded photoreceptors illustrating the measurements of tubulin width at three locations relative to the POC5 signal. 0: distal end of the CC-IS; +150: 150 nm distally to the CC-IS end; -150: 150 nm proximally to the CC-IS end. Scale bar: 500 nm. (**B**) Tubulin width measurements of the photoreceptor at the three locations depicted in (A) (−150 nm, 0 nm, +150 nm) from P4 to P60. ≥ 3 animals per time point. Scale bar: 200 nm. (**C**) Scheme describing the measurements of CC-IS signal end (green) and MTDs spread signal start used in (D). (**D**) Comparison of the position of the CC-IS end (POC5) or microtubule spread start relative to the centriole’s proximal end, from P4 to P60. ≥3 animals per time point. Note that IS measurements corresponds to the POC5 data presented in Fig. 2F. (**E**) EM transversal sections at the CC (top), or at the bulge (bottom). Dashed lines represent MTDs measured in F and G. Filled green arrowhead points to the CC-IS, empty green arrowhead reveals the absence of the CC-IS. Scale bar: 200 nm. (**F, G**) Distribution of the perimeter (F) or circularity (G) of the MTDs from transversal sections of photoreceptor CC (grey) or bulge (orange). N=1 animal. Means and standard deviations are listed in Table S1.

Finally, we tested whether the tubulin spread observed in some retinal degenerative diseases such as RP, is linked to the structural loss of the CC-IS and whether it precedes photoreceptor degeneration. For this purpose, we next took advantage of a RP28 mouse model deficient for FAM161A (*Fam161a*^*tm1b/tm1b*^), displaying progressive photoreceptor degeneration accompanied by loss of visual function from one month of age and associated with spread MTDs (*12*).

First, using tubulin and RHODOPSIN, we assessed rod OS development from P4 to P60 in *Fam161a*^*tm1b/tm1b*^ retinas and found no noticeable defects between P4 to P14, suggesting that early development of OS is not impacted (**Fig. 4A, Fig. S5A**). In contrast, from P30 onwards, RHODOPSIN signal progressively decreased, correlating with membrane discs disorganization and progressive loss of visual acuity in these animals (*12*). Consistently, tubulin signal in P60 reveals important defects with spread MTDs, recapitulating the phenotype previously seen in EM (*9*) (**Fig. 4A-D**). Similarly, we found that cone OPSIN, a marker of the cone photoreceptor cells, was also strongly disorganized in *Fam161a*^*tm1b/tm1b*^ retinas, with spread MTDs, indicating that cones are also degenerating in this model system (**Fig. S5B**). As *Fam161a*^*tm1b/tm1b*^ rods seem to develop normally until P14, we monitored FAM161A by immunostaining to evidence potential residual expression (**Fig. S6A, C**). We found that a faint signal of FAM161A was still observable at the level of the CC in early time points with ∼50% of CC positive until P14, suggesting that a truncated protein is still weakly expressed at early time points (*12*) (**Fig S6A, C**). In addition, we noticed that FAM161A seemed more stable at centrioles in *Fam161a*^*tm1b/tm1b*^ mice, before vanishing with solely 12% of FAM161A positive centrioles at P30 (**Fig. S6A, C**). We further validated this pattern of residual FAM161A signal in a second mouse model deficient for FAM161A (*Fam161a*^*GT/GT*^), known to generate a truncated FAM161A protein (*9*) (**Fig S6B**).

**Fig. 4.**
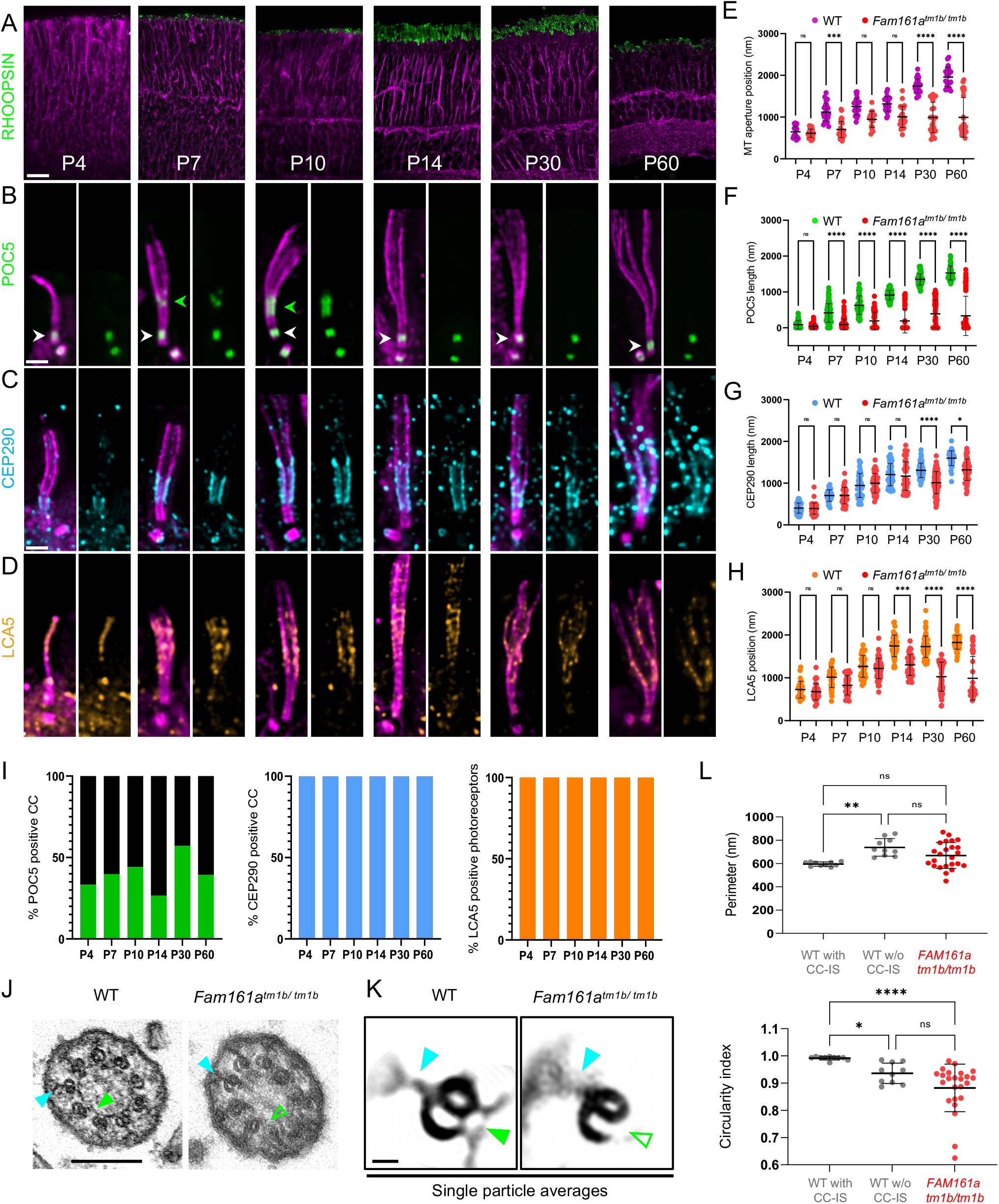
FAM161A mutation leads to the specific loss of the CC-inner scaffold. (**A**) Low magnification of expanded *Fam161a*^*tm1b/tm1b*^ retinas stained for RHODOPSIN (green) and tubulin (magenta) from P4 to P60. Scale bar: 20 µm. (**B-D**) Expanded *Fam161a*^*tm1b/tm1b*^ photoreceptors stained for POC5 (green, B), CEP290 (cyan, C) and LCA5 (dark yellow, D) from P4 to P60. Note the transitory appearance of the CC-IS between P7 and P10 (green arrowheads) followed by its collapse, paralleling microtubule spread. In contrast, centriole IS (white arrowheads) is kept over time. Scale bar: 500 nm. (**E**) Comparison of the microtubule enlargement position relative to MC proximal end between WT (green) and *Fam161a*^*tm1b/tm1b*^ (red) photoreceptors. ≥3 animals per time point. (**F-H**) Impact of the *Fam161a*^*tm1b/tm1b*^ mutant on CC-IS length (POC5 staining, F), CEP290 length (G) or LCA5 start position (H). ≥3 animals per time point. (**I**) Percentage of CC positive for POC5 (green), CEP290 (cyan) and photoreceptor positive for LCA5 (dark yellow) from P4 to P60 in *Fam161a*^*tm1b/tm1b*^ photoreceptors. ≥3 animals per time point. (**J**) EM micrographs of WT (left) or *Fam161a*^*tm1b/tm1b*^ (right) CC transversal sections. Scale bar: 200 nm. (**K**) single particle averages of WT or *Fam161a*^*tm1b/tm1b*^ MTDs. Note that Y-links are present in both conditions (cyan arrowheads) whereas CC-IS is present in WT (full green arrowhead) but absent in *Fam161a*^*tm1b/tm1b*^ (empty green arrowhead). Scale bar: 20 nm. (**L**) Comparison of the perimeter or the circularity of the microtubule axoneme from transversal sections of WT (grey) or *Fam161a*^*tm1b/tm1b*^ (red) photoreceptors. Note that WT data are the same used in Fig. 3I, J. N= 1 animal. Note that in panels E-H and K, WT measurements are the same as presented in Fig. 2E-G and Fig. 3F-G. Means, percentage and standard deviations are listed in Table S1.

We next monitored the localization of POC5 and CENTRIN during the development of photoreceptors in *Fam161a*^*tm1b/tm1b*^ or *Fam161a*^*GT/GT*^ retinas (**Fig. 4B**, and **Fig. S6A**). Similarly to FAM161A, POC5 and CENTRIN proteins showed an early and transitory localization at the CC before disappearing later on in the majority of the photoreceptors observed, despite mutant animals being heterogeneously impacted over time (**Fig. 4B, F, I** and **Fig. S6**). This loss is paralleled with an opening of the axonemal MTDs, that progressively reaches the mother centriole distal end, indicative of a shortening of the CC region as previously described (*9*) (**Fig. 4B, E, F**). This MTDs spread becomes prominent at P30, where OS layer starts to be impacted, accounting for the beginning of visual function decline at this age (*12*). In contrast to FAM161A, we noticed that POC5 and CENTRIN remain present at centrioles all over the analyzed time points, suggesting a differential impact of FAM161A mutation on CC- and centriole-IS structures, that could explain the exclusive retinal phenotype observed in RP28 (**Fig. 4B**, and **Fig. S6**). To further understand this result, we monitored POC5 in human U2OS cycling cells depleted of FAM161A using siRNAs. While FAM161A was absent in 89% of centrioles in FAM161A-depleted cells, only 38% of centrioles had lost POC5 (**Fig. S7A-D**), suggesting that FAM161A does not control POC5 recruitment at centrioles, consistently to the results obtained in photoreceptors. In contrast, we found that FAM161A was reduced similarly to POC5 at centrioles in POC5-depleted U2OS cells (**Fig. S7A-D**), unveiling that POC5 mostly controls FAM161A localization at centrioles. Finally, co-depletion of both proteins strongly affects FAM161A and POC5 presence at centrioles and induces in ∼15% of cases centriole architectural defects with broken microtubule pieces, a phenotype reminiscent to the spread MTDs in the connecting cilium in FAM161A mutants (**Fig. S7E**).

We also investigated whether CEP290 and LCA5 are affected in *Fam161a*^*tm1b/tm1b*^ photoreceptors (**Fig. 4C, D**). In contrast to POC5, 100% of photoreceptors displayed CEP290 or LCA5 signals over the analyzed time points, reinforcing the independence of these different modules (**Fig. 4I**). We observed a decrease in CEP290 length from P30 onwards, presumably as a secondary consequence of MTDs defects affecting axoneme integrity at this time point (**Fig. 4C, G**). LCA5 localization was also not impaired at early time points, but then re-localized closer to the mother centriole from P14, consistent with the shortening of the CC and revealing the disorganization of the bulge region, possibly causing membrane discs formation impairment (**Fig 4D, H**).

Altogether, these results show that a specific loss of CC-IS proteins, which initially localize transitorily in *Fam161a*^*tm1b/tm1b*^ retinas and presumably allow early OS formation, triggers MTDs destabilization and CC shortening, prior to OS degeneration. We next investigated whether this CC-IS proteins loss is correlated with a loss of the CC-IS structure by analyzing transversal EM sections of a *Fam161a*^*tm1b/tm1b*^ P30 mouse retina (**Fig. 4J** and **Fig. S8**). We found that the vast majority of the axonemal sections lacked CC-IS, whereas Y-links were still present, confirming the molecular architecture findings by U-ExM. Moreover, the global MTDs structure was sometimes greatly impaired with opened B-microtubules and with membrane invaginations found inside the axoneme (**Fig. S8**). Consistently, axonemal perimeter distribution was largely heterogeneous in mutant photoreceptors, together with a less round shape, demonstrating that the lack of the CC-IS impairs MTDs cohesion and OS organization (**Fig. 4J, L**). To ascertain this lack of CC-IS, we performed particle averaging on WT and *Fam161a*^*tm1b/tm1b*^ MTDs from EM micrographs and highlighted the underlying structural defects (**Fig. 4K**, lower panel, **Fig. S9**). This analysis confirmed the presence of the Y-links as well as the complete structural loss of the CC-IS in *Fam161a*^*tm1b/tm1b*^.

Finally, we studied the effect of the CC-IS loss on overall photoreceptor cell integrity and in particular the rod outer segment membrane stacks containing RHODOPSIN, essential for proper vision (**Fig. 5 A-D**). We found that while RHODOPSIN is correctly located above the CC in P60 WT cells, the loss of CC-IS induces a collapse of the OS structure, with the RHODOPSIN signal and membrane discs displaced towards and inside the CC region (**Fig. 5B, D** and **Fig. S8**). Moreover, in addition to a mislocalization of RHODOPSIN at the cell body as previously observed (*9*) (**Fig. S10**, white arrows), we noticed in several cells an accumulation at the base of the CC, potentially indicating a defect in RHODOPSIN transport in *Fam161a*^*tm1b/tm1b*^. These results demonstrate that CC-IS acts as a structural foundation to maintain the cell architecture and a correct outer segment organization.

**Fig. 5.**
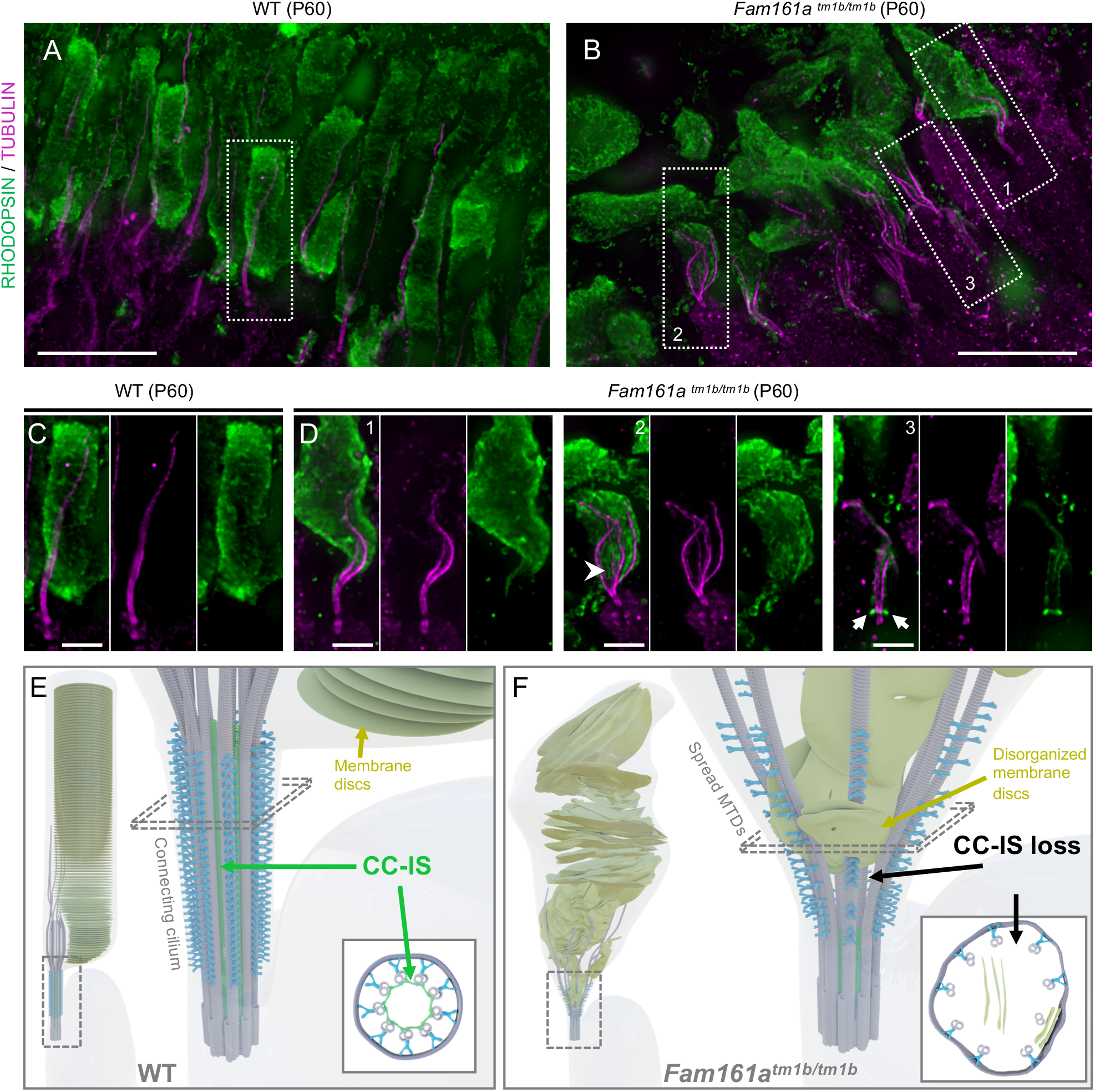
Loss of the CC-inner scaffold causes photoreceptor degeneration. (**A, B**) Low magnification of expanded WT (A) or *Fam161a*^*tm1b/tm1b*^ (B) retinas stained for RHODOPSIN (green) and tubulin (magenta) at P60. Note the microtubule defects in the mutant, accompanied by outer segment collapse. Dashed lines represent the insets (1, 2 and 3) depicted in C, D. Scale bar: 20 µm. (**C-D**) Insets from WT (C) or *Fam161a*^*tm1b/tm1b*^ (D) retinas depicted in (A) and (B), respectively. Arrowhead shows the RHODOPSIN signal entering inside the microtubules. Arrows depicts the accumulation of the RHODOPSIN at the base of the CC. Scale bar: 5 µm. (**E, F**) Model representing mature WT (E) or Fam161atm1b/tm1b (F) photoreceptor with a highlight on the connecting cilium region. Schemes representing transversal sections are depicted in the bottom rights of each panel.

In summary, our work highlights the presence of an inner scaffold structure inside the connecting cilium, containing FAM161A, POC5 and CENTRIN, that is crucial for proper maintenance of the photoreceptor OS in mouse (**Fig. 5E, F**). Moreover, this work provides a molecular and structural basis to understand photoreceptor degeneration in *retinitis pigmentosa* linked to ciliary defect, paving the way for future therapeutic options to restore visual acuity in RP patients.

## Acknowledgments

We warmly thank Dror Sharon, Avigail Beryozkin and Thomas Langmann for sharing their FAM161A-deficient mouse strains and Susanne Borgers, Sylvie Montessuit and Jean-Claude Martinou for technical help. We thank the PFMU (UniGE) for EM samples preparation, the Bioimaging center (UniGE), and Marine. H. Laporte for providing the centrioles measurements in Fig. S1. We also thank Gabriel Aeschlimann for the design of the model in Fig. 5.

## Author contributions

O.M. performed all the experiments described in the paper. A. G. and E. B. performed the siRNA experiments in human cells. Y. S. imaged and analyzed the EM micrographs. C.K., N.C. and Y.A. provided all mouse retinas (authorization VD1367). V.H. and P.G. conceived, designed and supervised the project with inputs from C. K., Y. A. and O. M. All authors wrote and revised the final manuscript;

## Data and materials availability

All data is available in the main text or the supplementary materials.

## Supplementary Materials

Materials and Methods

Figures S1-S10

Tables S1-S2

