## Supplemental data for "The connecting cilium inner scaffold provides a structural foundation to maintain photoreceptor integrity"

### Supplementary Materials

**Materials and Methods**

Mouse model

Animals were handled in accordance with the statement of the "Animals in Research Committee" of the Association for Research in Vision and Ophthalmology, and experiments were approved by the local institutional committee (VD1367). The mice were maintained at 22 °C with a 12-h light/12-h dark cycle with light on at 7:00 AM and were feed ad libitum. *Fam161A*-deficient mice, *Fam161a*$^{tm1b/tm1b}$ (*1*) and *Fam161a*$^{GT/GT}$ (*2*), were obtained from Avigail Beryozkin and Thomas Langmann, respectively. C57BL6RJ (Janvier Labs, Le Genest-Saint-Isle, France) were used as controls.

Mice were sacrificed at postnatal day P4, P7, P10, P14, P30 (1 month) and P60 (2 months) of age and eyes enucleated.

Retina dissection

After enucleation, eyes were fixed for 15 min at room temperature in 4% PFA (paraformaldehyde, P6148, Sigma-Aldrich) in phosphate-buffered saline (PBS) (P4, P7, P10, P14) or 2% PFA in PBS (1 month and 2 months) and then transferred into (PBS). Then, cornea and lens were removed with micro scissors, and the sclera was separated from the retina. Retinas were then either kept as a cup for EM processing or incised to flatten it as a clover inside a 10 mm microwell of a 35 mm petri dish (P35G-1.5-10-C, MatTek) to allow their processing by Ultrastructure expansion microscopy (U-ExM).

Retina expansion

Protocol for retina expansion was adapted from the U-ExM method (*3*) with few optimizations. Briefly, crosslinking prevention step was extended to overnight (ON) incubation with 100 µL of 2% acrylamide (AA; A4058, Sigma-Aldrich) + 1.4% formaldehyde (FA; F8775, Sigma-Aldrich) at 37°C inside the 10 mm microwell of a petri dish (MatTek). Then, the solution was removed and 35 µL monomer solution (MS) composed of 25 µl of Sodium Acrylate (stock solution at 38% (w/w) diluted with nuclease-free water, 408220, Sigma-Aldrich), 12.5 µl of AA, 2.5 µl of N, N'-methylenbisacrylamide (BIS, 2%, M1533, Sigma-Aldrich) and 5 µl of 10X PBS was added for 90 min at RT to allow its penetration into the tissue. Then, MS was removed and 90 μL of MS was added together with ammonium persulfate (APS, 17874, ThermoFisher) and tetramethylethylendiamine (TEMED, 17919, ThermoFisher) as a final concentration of 0.5% for 45 min at 4°C first followed by 3h incubation at 37°C to allow gelation. A 24 mm coverslip was added on top to close the chamber. After, the coverslip was removed and 1 mL of denaturation buffer (200 mM SDS, 200 mM NaCl, 50 mM Tris Base in water, pH 9) was added into the MatTek dish for 15 min at RT with shaking. Then, careful detachment of the gel from the dish with a spatula was performed and the gel was incubated in 1,5 mL tube filled with denaturation buffer for 1h at 95°C and then ON at RT. The day after, the gel was cut around the retina that is still visible at this step, and expanded in three successive ddH2O baths. Then, the gel was sliced in order to have transversal sections of the retina that were then processed for immunostaining.

Human cell culture and expansion

Human U2OS were cultured similarly to (*4*). Basically, cells were grown in DMEM supplemented with GlutaMAX (Thermo Fisher Scientific), 10% fetal calf serum (Thermo Fisher Scientific), and penicillin and streptomycin (100 μg/ml, Thermo Fisher Scientific). Expansion on U2OS cells was performed as previously described (*5*).

Immunostaining

After expansion, human cell gels were shrinked with three 5-min baths of 1x PBS. Then primary antibodies were incubated for 3h at 37°C in PBS with 2% of Bovine Serum Albumine (BSA). Gels were washed 3 times 10 min in PBS with 0.1% Tween 20 (PBST) prior to secondary antibodies incubation for 3h at 37°C. After a second round of washing step (3 times 10 min in PBST), gels were expanded with 3 baths of ddH20 before imaging. Antibodies used are referenced in Table S2. Tubulin staining presented in the figures corresponds to either a mixture of anti α- and β-tubulin (raised in mouse) or anti α-tubulin alone (raised in rabbit) depending on the species of the other protein stained.

For retina slices immunostaining, primary antibodies incubation was prolonged ON at 4°C. Image acquisition was performed on an inverted confocal Leica TCS SP8 microscope or on a Leica Thunder DMi8 microscope using a 20x (0.40 NA) or 63 ×

(1.4 NA) oil objective with Lightening or Thunder SVCC (small volume computational clearing) mode at max resolution, adaptive as 'Strategy' and water as 'Mounting medium' to generate deconvolved images. 3D stacks were acquired with 0.12 μm z-intervals and an x, y pixel size of 35 nm.

Human cell transfection

U2OS cells were plated onto coverslips in a 6-well plate at 100'000 cells/well 24h before transfection. Using Lipofectamine RNAimax (Thermo Fischer Scientific), cells were transfected either with 50 nM of silencer select negative control siRNA1 (4390843, Thermo Fisher), 25 nM of siRNA against POC5, or a mixture of 12.5 nM of two different siRNAs against FAM161A. For double transfections, a mixture of 25 nM Poc5 siRNA with both FAM161A siRNAs at 12.5 nM each was used. Medium was changed 6h post-transfection and cells were analyzed 72h after transfection. siRNA references and sequences are referenced in Table S2.

Electron Microscopy

For electron microscopy (EM) analyses, retina cups were first incubated ON at RT with 3% PFA (15710, Electron Microscopy Sciences) and 0,1% glutaraldehyde (16200, Electron Microscopy Science) in PBS. Samples were further treated with 2% osmium tetraoxyde (05500-1g, Sigma-Aldrich) in buffer for 30 minutes and immersed in a solution of uranyl acetate (21447-25g, Polysciences) 0.25% ON to enhance contrast of membranes. Samples were dehydrated in increasing concentrations of ethanol followed by pure propylene oxide (82320-1L, Sigma-Aldrich), and then embedded in Epon resin. Serial ultra-thin sections of 50 or 100 nm were finally cut and stained with 5% uranyl acetate (in H2O) and Reynolds' lead citrate (*6*). Micrographs of the WT sample were acquired using a G2 Sphera microscope operated at 120 kV equipped with an Eagle detector at a magnification of 25000x corresponding to a pixel size 4,5 A. The micrographs of the mutant sample were acquired using a Talos LC120 microscope operated at 120 kV equipped with a CetaD detector at a magnification of 36000x corresponding to pixel size 4 A. All micrographs were acquired with a defocus between -3 to -5 μm.

Symmetrization

Symmetrization of EM micrographs was done using CentrioleJ plugin (*7*). The first step consists in the circularization of the pattern (here the CC), to correct elliptical deformation of the acquisition, by manually picking all the center of mass of the A-microtubules. Then, symmetrization consists in rotating the source image according to its symmetry (here 9-fold) and sum-projecting all the rotated images.

Particle classification and averaging

745 and 178 particles were manually picked on the microtubule doublets and extracted with box size of 310 and 256 pixels from wild-type and mutant, respectively. The particles were subjected to 2D classification procedure implemented in Cryosparc 3.2 (*8*). The alignment resolution and maximum resolution were limited to 50 and 10 Å. respectively for alignment over 40 iterations. Moreover, Force Max over poses/shifts, Enforce non-negativity and Use clamp-solvent to solve 2D classes were set to true to reduce background noise.

Quantifications

*Expansion Factor*

The expansion factor of each experience was calculated in a semi-automated way by comparing the Full Width at Half Maximum (FWHM) of photoreceptor mother centriole proximal tubulin signal with the proximal tubulin signal of expanded human U2OS cell centrioles using PickCentrioleDim plugin described elsewhere (*9*). Briefly, for each experiment, at least 10 photoreceptor mother centrioles FWHM were measured and compared to a pre-assessed value of U20S centriole width (25 centrioles: mean= 231.3 nm +/-15.6 nm). The ratio between measured FWHM and known centriole width gave the expansion factor.

*Protein shift*

To calculate the protein shift compared to tubulin, confocal top view images of connecting cilia were analyzed. Using Image J, a line crossing connecting cilia on their diameter was drawn and plot profiles of each channel (protein of interest and tubulin) were generated. Then, distances between peak intensities of the protein of interest and tubulin were measured and corrected with the mean expansion factor calculated from all experiments (Fig S1D). For each connecting cilium, 4 measures were done to correct potential tilting effects.

*Protein signal length and position*

Protein signal length or position compared to mother centriole proximal end (depicted with tubulin) were measured using a segmented line drawn by hand (ImageJ) to fit with photoreceptor curvature, and corrected with the expansion factor. Only photoreceptors where both protein signal (POC5, CEP290 or LCA5) and centriole proximal end (tubulin) were clearly visible were selected for measurements.

Tubulin *spread*

Tubulin spread was assessed at each time point in a semi-automated way by measuring FWHM of tubulin signal with PickCentrioleDim plugin (*9*) on three different locations of the photoreceptor corresponding to 150 nm proximally to the end of the CC POC5 staining (-150), at the level of the end of the CC POC5 staining (0), or 150 nm distally to the end of the CC POC5 staining (+150). Each measurement was subsequently corrected with its respective expansion factor. The position of the microtubule opening was measured manually using a segmented line drawn in ImageJ (*10*), to fit with photoreceptor curvature, and corrected with the expansion factor. Independent tubulin staining alone was used to avoid bias with CC-IS staining.

*Microtubule axoneme perimeter and circularity*

Microtubule axoneme perimeter and circularity were assessed from EM images. Using ImageJ, elliptical lines were drawn by hand to fit with microtubule doublets center allowing the measurements of the perimeter and the area. Circularity (Ci) was calculated using the formula: $Ci = 4*Pi*Area / Perimeter^2$. Only axonemes with all the microtubule doublets visible were quantified. Bulge regions were defined as lacking the inner scaffold ring and the Y-link structures.

*siRNA*

In order to evaluate siRNA efficiency after transfection, POC5 and FAM161A stainings after U-ExM were first assessed by eyes by measuring the proportion of centrosomes (mother and daughter centrioles) with a complete signal at both centrioles (non-depleted) and centrosomes with one or both centrioles with a reduced or absent signal for the proteins (depleted).

For fluorescence intensity measurements of FAM161A and POC5, maximal projections were used using Fiji (*10*) on non-deconvoluted images. The same circular Region Of Interest (ROI) drawn by hand was used to measure protein signal intensities around every centriole and their corresponding background. Fluorescence intensity was finally calculated by subtracting the average value of two background measures (raw integrated density).

Statistical analyses

The comparison of two groups was performed using non-parametric Mann-Whitney test, if normality was not granted because rejected by Pearson test. The comparisons of more than two groups were made using non-parametric Kruskal-Wallis test followed by post-hoc test (Dunn's for multiple comparisons) to identify all the significant group differences. Every measurement was performed on at least three different animals, or cell culture independently, unless specified. Data are all represented as a scatter dot plot with centerline as mean, except for percentages quantifications, which are represented as histogram bars. The graphs with error bars indicate SD (+/-) and the significance level is denoted as usual (*$p<0.05$, **$p<0.01$, ***$p<0.001$, ****$p<0.0001$). All the statistical analyses were performed using Prism9. Every mean, percentage, standard deviation and tests used for comparison are referenced in Table S1.

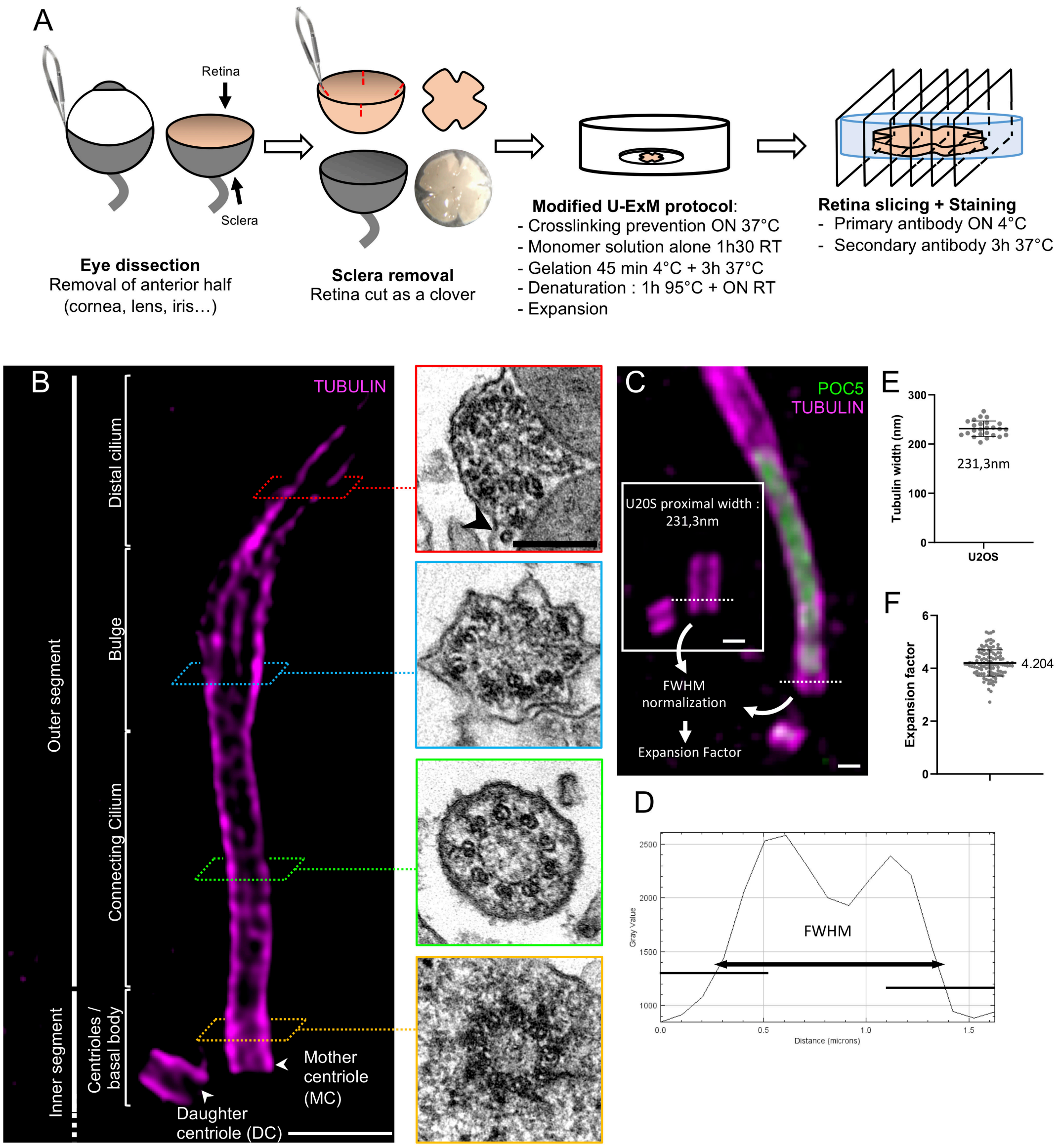


**Fig. S1. U-ExM protocol adapted to mouse retina**

(**A**) Scheme resuming the main steps of mouse retina dissection and expansion. (**B**) Expanded photoreceptor (left) highlighting the different regions of the distal inner segment and outer segment thanks to the tubulin staining, together with the corresponding EM images (right). Scale bars: left= 500 nm; right= 200 nm. Black arrowhead points to microtubule singlet in the distal cilium. Note that the EM picture depicting the bulge region is the same as in the main figure. (**C**) Photoreceptor expansion factor calculation by comparing tubulin signal width at the photoreceptor mother centriole's proximal end with the tubulin width of U2OS centriolar proximal end. Scale bars: 200 nm. (**D**) Fluorescence plot profile of tubulin at the level of the photoreceptor mother centriole proximal end generated with PickCentrioleDim plugin. FWHM: Full Width at Half Maximum. (**E**) Measurements of U2OS centriole proximal width. N=4 independent experiments. See Table S1. (**F**) Distribution and mean of the expansion factors calculated in all the experiments. > 40 animals were used to measure expansion factor. See Table S1.

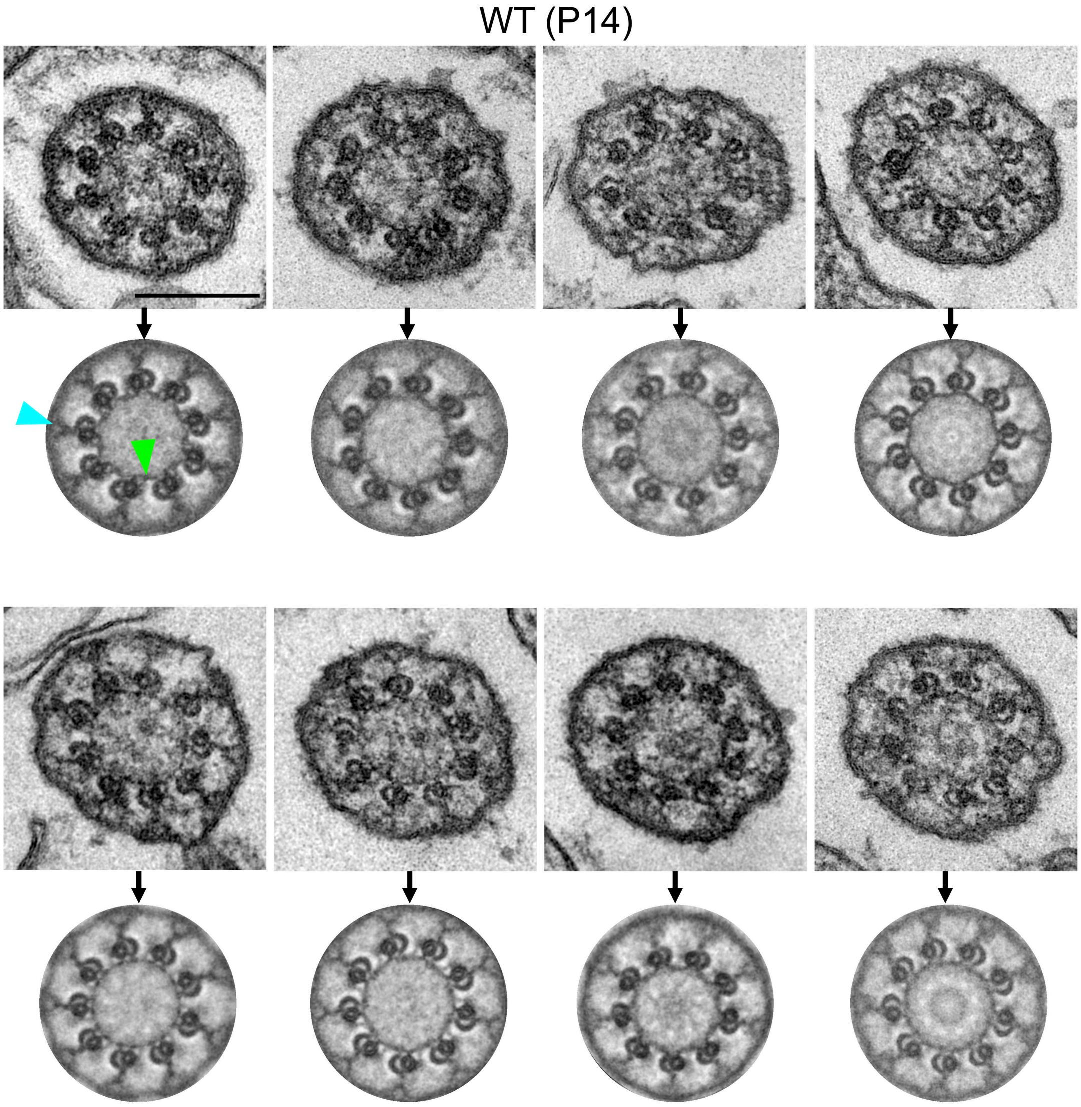


**Fig. S2. Gallery of raw and symmetrized EM images of P14 WT CC transversal sections.**

EM micrographs of WT P14 connecting cilia before and after symmetrization using CentrioleJ (see methods), highlighting the presence of the CC-IS (green arrowhead) and the Y-links (blue arrowhead). Scale bar: 200 nm

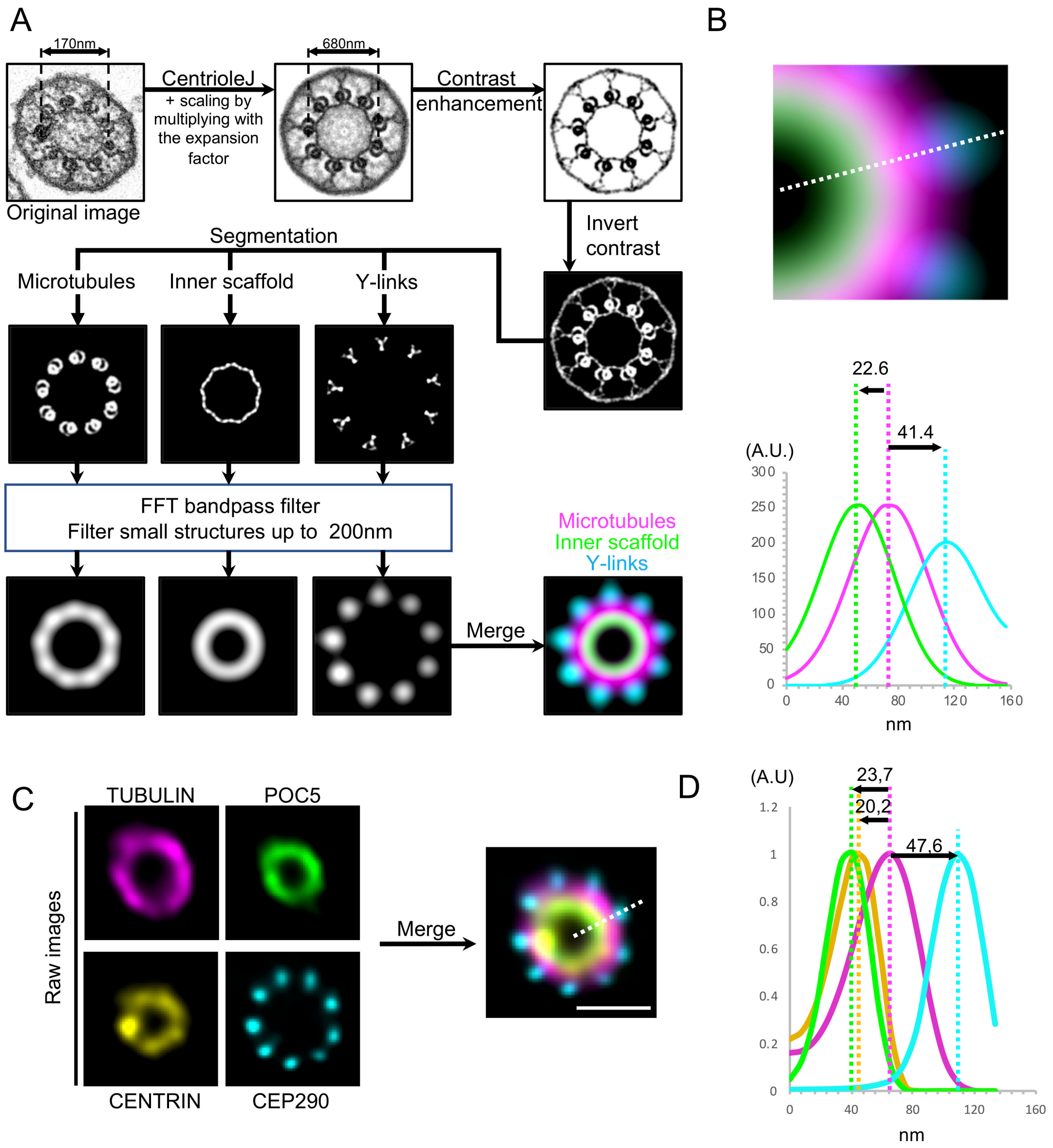


**Fig. S3. Simulation of U-ExM signals from EM micrographs**

**(A)** Scheme representing the simulation pipeline as previously used in (*3*). First, a raw EM micrograph was symmetrized using CentrioleJ (see Methods). Then, after contrast optimization, all the different structures (microtubules, inner scaffold and Y-links) were segmented and submitted to a bandpass filter mimicking the limit of resolution obtained with classic fluorescence microscopy (200 nm). The simulated signals of each structure were then merged to reconstruct the final simulation. **(B)** Relative distance of each structure based on the reconstructed simulation. Top: representation of the line (white dashed) drawn to make the measurement of peak intensities for each structure (inner scaffold in green, microtubules in magenta, and Y-links in cyan). Bottom: peak intensity distances of each structure. (**C**) Merge of independent stainings of POC5 (green), CENTRIN (yellow) and CEP290 (Cyan) to compare measurements with the simulation. The line drawn to make the measurement of peak intensities for each staining is represented with the white dashed line. Scale bar: 200 nm. **(D)** Peak intensity distances of each protein calculated from the merge image in (C).

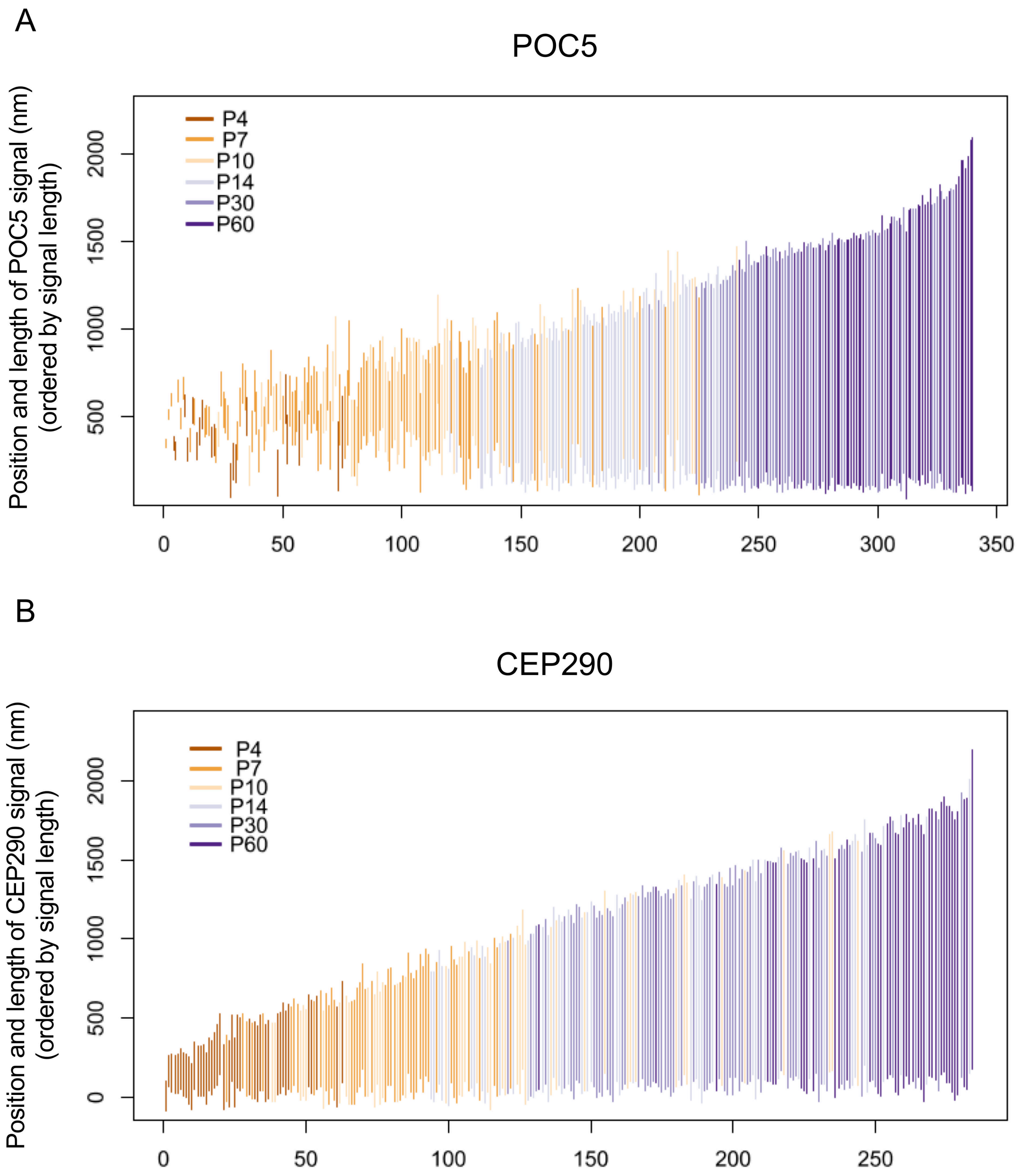


**Fig. S4. CC POC5 and CEP290 growth over time**
Graphs representing the position and the length of the POC5 (A) and CEP290 (B) CC signals sorted by length. Each color depicts the age of the animals measured, confirming that the temporality of the growth of the two signals.

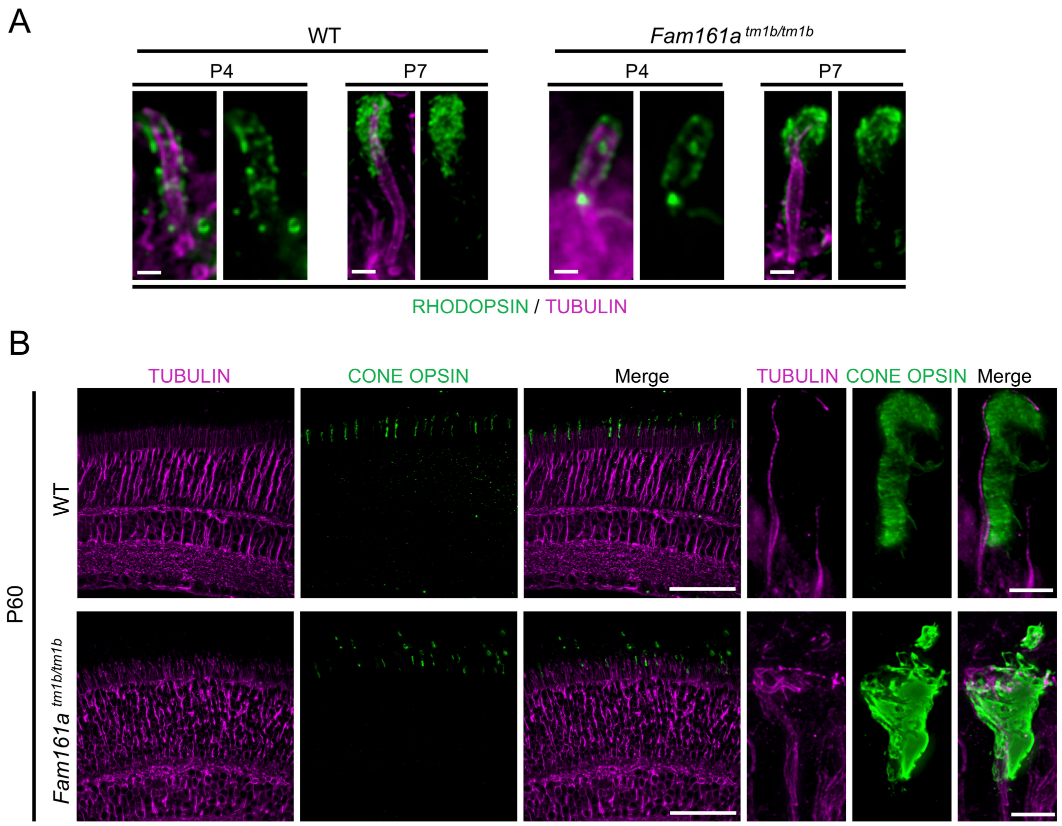


**Fig. S5. Impact of Fam161a$^{tm1b/tm1b}$ on RHODOPSIN and CONE OPSIN staining over time**.

(**A**) Comparison of early outer segment development in WT and Fam161a$^{tm1b/tm1b}$ photoreceptors. Note that RHODOPSIN outlines the outer segment tubulin signal at P4 and then accumulates distally at P7. Scale bar: 500 nm. (**B**) Expanded P60 WT or *Fam161a$^{tm1b/tm1b}$* retinas stained for CONE OPSIN (green) and tubulin (magenta). High magnification of the cone photoreceptors (right) shows that cones axonemes are also greatly impacted in *Fam161a$^{tm1b/tm1b}$* retinas, revealing no obvious difference between cones and rod at P60 in mutant mice. Scale bar: low mag= 50 µm; High mag= 2 µm.

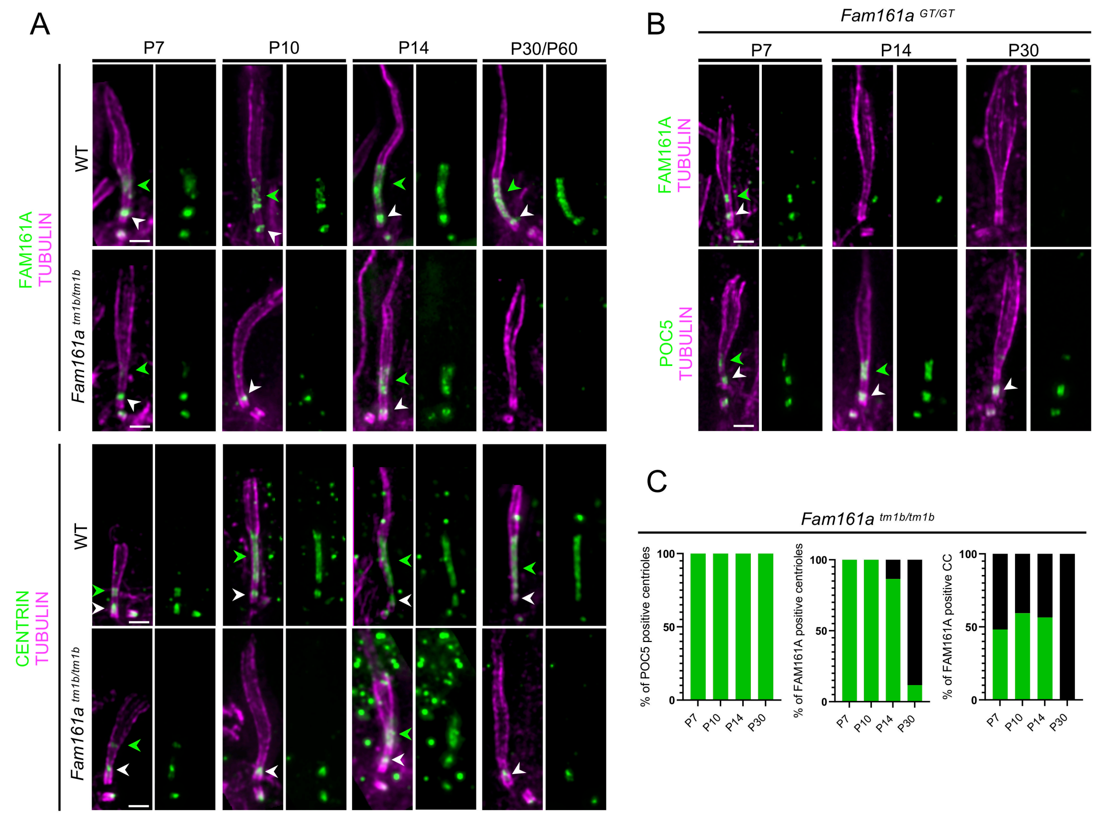


#### Fig. S6. FAM161A and CENTRIN staining in the two mouse models of RP28

(**A**) Expanded WT and *Fam161a*$^{tm1b/tm1b}$ photoreceptors stained for tubulin and FAM161A (top) or CENTRIN (bottom) at different ages. Note that P30/P60 represents a mixture of P30 or P60 images. Scale bar: 500 nm. White arrowheads point to centriole IS and green arrowheads point to CC-IS, when present. (**B**) Expanded *Fam161a*$^{GT/GT}$ photoreceptors stained for tubulin and FAM161A (top) or POC5 (bottom) between P7 and P30. Scale bar: 500 nm. White arrowheads point to centriole IS and green arrowheads point to CC-IS, when present. (**C**) Proportion of POC5-positive centrioles (left), FAM161A-positive centrioles (center) or FAM161A-positive CC (right) in *Fam161a*$^{tm1b/tm1b}$ photoreceptors between P7 and P30.

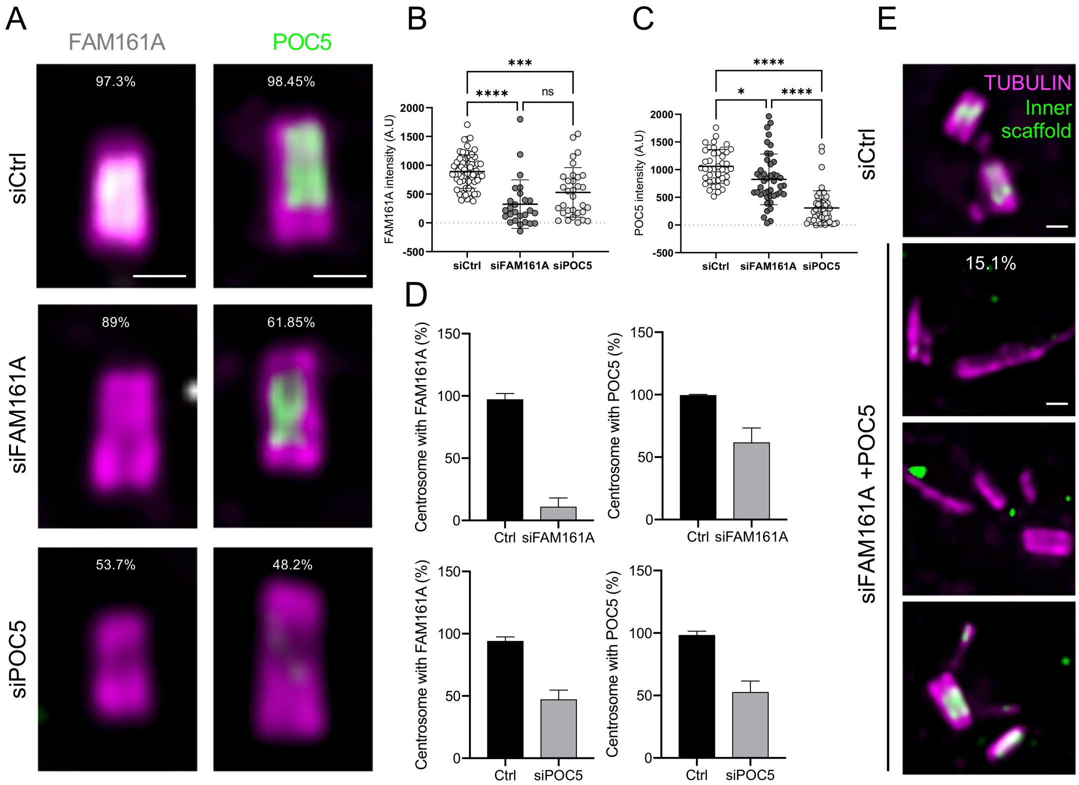


**Fig. S7. Impact of POC5 or FAM161A siRNA in human centrioles**

(**A**) Representative widefield images of expanded U2OS centrioles treated with siCtrl, siPOC5 or siFAM161A stained with tubulin (magenta) and FAM161A (gray) or POC5 (green). Scale bar: 200 nm. (**B**) Mean fluorescence intensity of FAM161A in indicated conditions. (**C**) Mean fluorescence intensity of POC5 in indicated conditions. (**D**) Percentage of positive centrosomes (with FAM161A or POC5 staining) in siCtrl, siPOC5 or siFAM161A treated cells. (**E**) Representative widefield images of expanded U2OS centrioles treated with siCtrl or siFAM161A+ siPOC5 stained with tubulin (magenta) and inner scaffold protein (green). Scale bar: 200 nm. ≥3 independent experiments for each measurement.

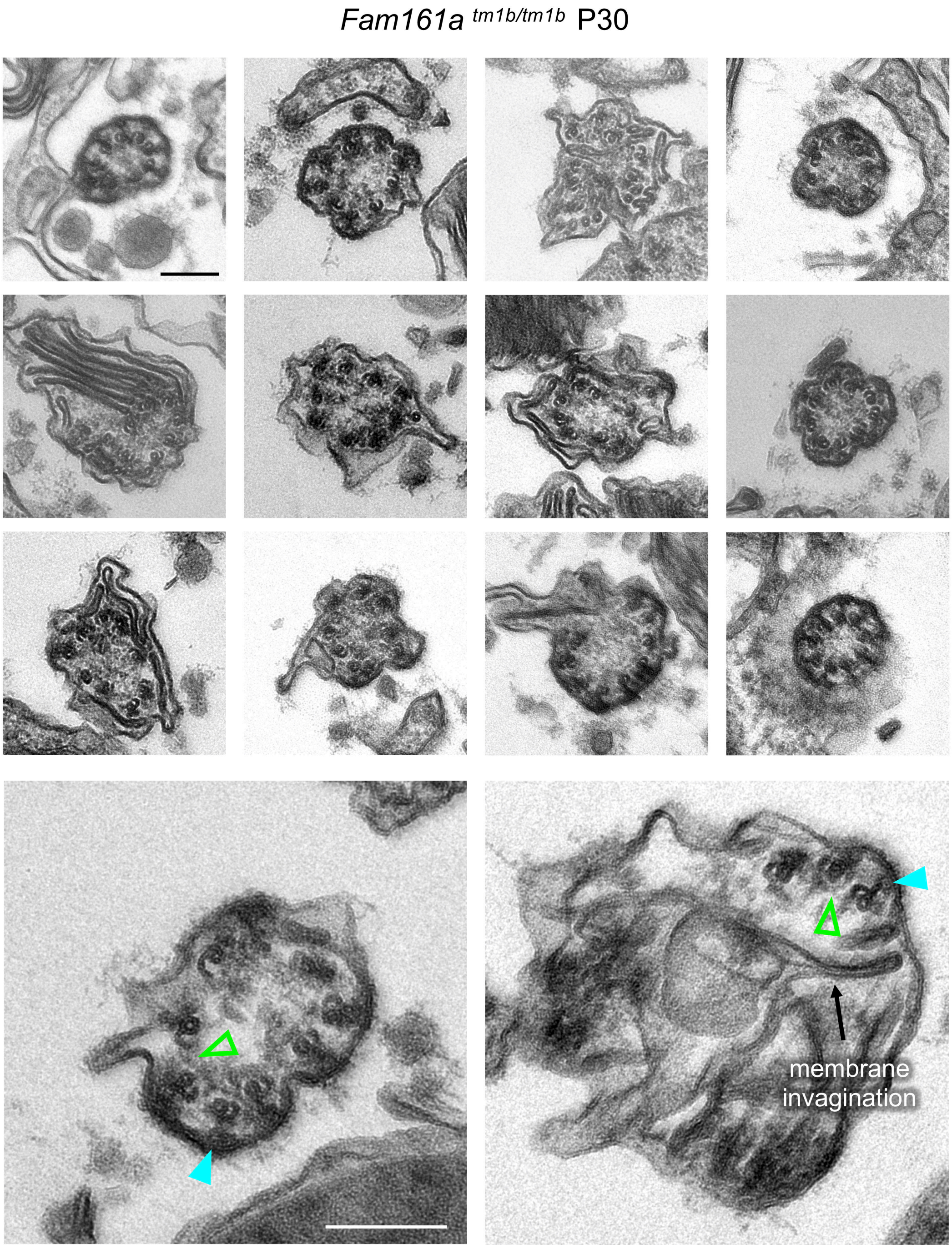


**Fig. S8. EM gallery of P30 *Fam161a*$^{tm1b/tm1b}$ connecting cilium transversal sections.**

EM micrographs of mutant connecting cilia revealing the loss of cohesion of MTDs within the axoneme. Note that Y-links are still observable in some MTDs, even in strongly affected CC where membrane invaginations are present within the axoneme (bottom right). Empty green arrowheads indicate the lack of CC-IS. Blue arrowhead highlights the presence of Y-links. Scale bars: 200 nm

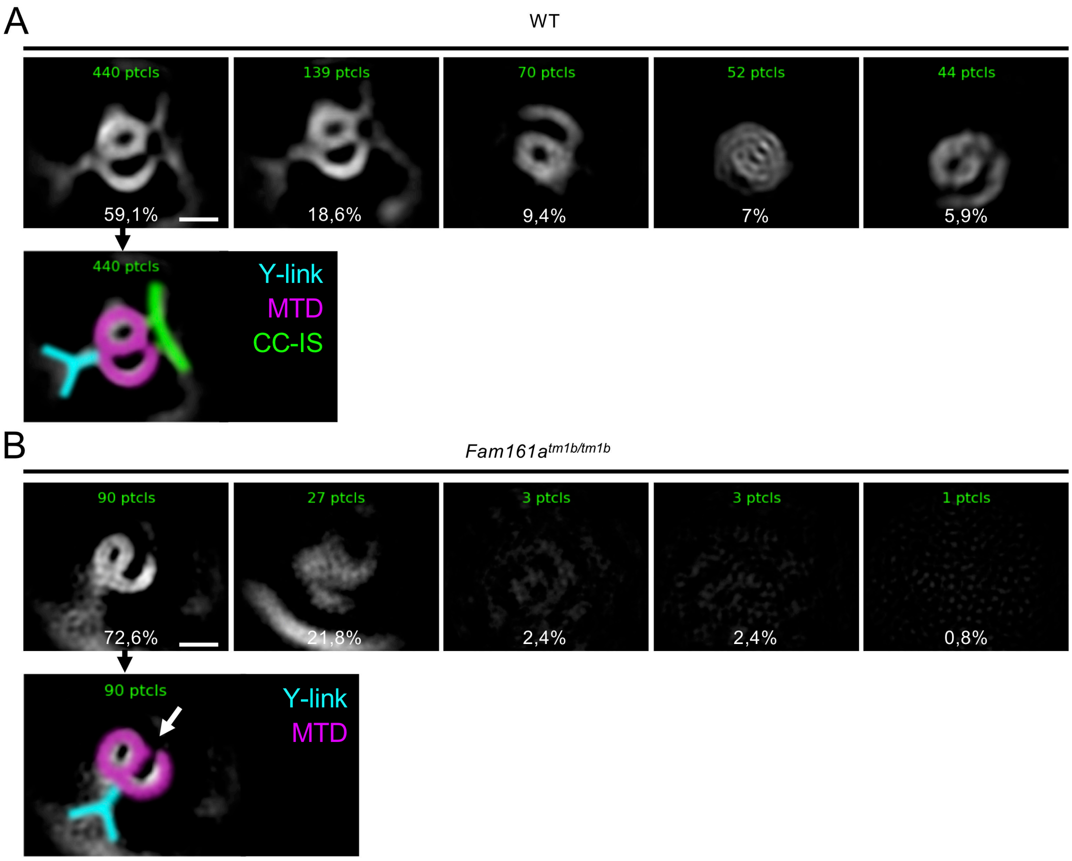


**Fig. S9. Single particle average classification of WT or *Fam161a*$^{tm1b/tm1b}$ microtubule doublets.**

Representation of the classification (5 classes) of the particle averaging obtained from WT (A) or *Fam161a*$^{tm1b/tm1b}$ (B) microtubule doublets (See Methods). The number of particles in each class is written in green, and the relative representation of each class is depicted below. For the most representative class of either WT or *Fam161a*$^{tm1b/tm1b}$, superimposition of the different structures (Microtubules in magenta, Y-links in Cyan and the CC-IS in green) was drawn for illustration purposes. Scale bar: 20 nm.

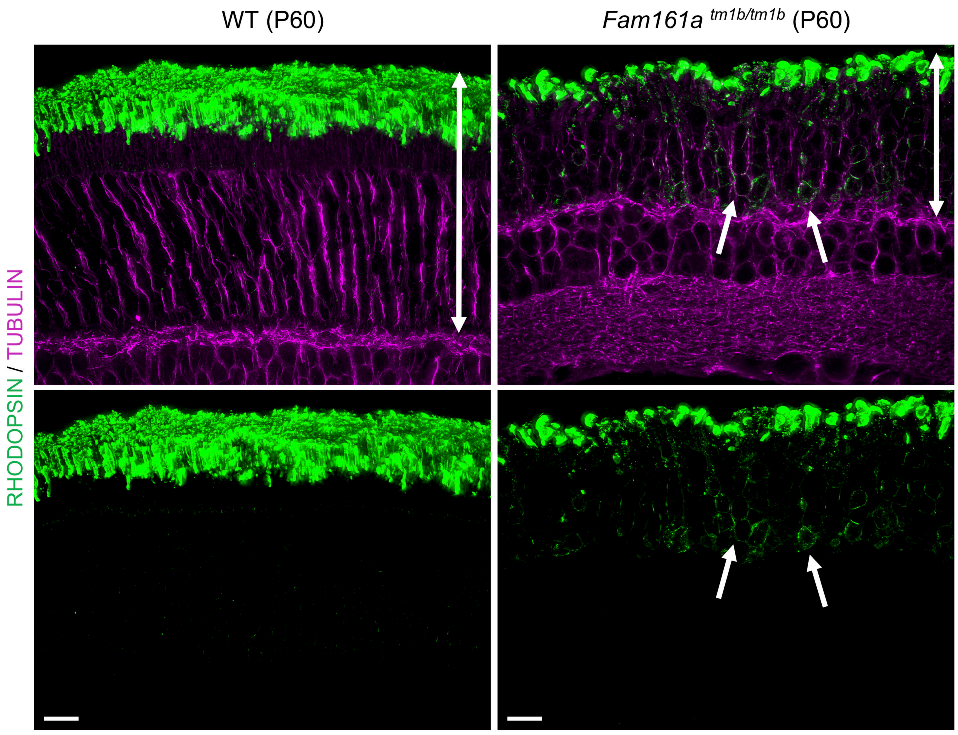


**Fig. S10. RHODOPSIN mis-localization in late *Fam161a*$^{tm1b/tm1b}$ retinas**

(**A**) Expanded P60 WT or *Fam161a*$^{tm1b/tm1b}$ retinas stained for RHODOPSIN (green) and tubulin (magenta). White arrows show RHODOPSIN signal at the level of photoreceptor cell bodies, notably around the nuclei in *Fam161a*$^{tm1b/tm1b}$ retinas. Double headed arrows reveal the difference of photoreceptor layer thickness between WT and *Fam161a*$^{tm1b/tm1b}$. Scalebar: 50 µm

**Table S1: Descriptive statistics for all measurements**

**Table S2: Antibodies and siRNA references**
